# PLANCK: super-multiplex optical imaging without labeling

**DOI:** 10.64898/2026.07.02.736216

**Authors:** Xinwen Liu, Xuemeng Li, Lele Xu, Mian Wei, Areej Niaz, Ye He, Wei Min

## Abstract

Molecular information is vital for imaging technology. Optical imaging acquires molecular specificity almost exclusively via labeling strategy, which is fundamentally constrained by limited multiplexing capacity, high running costs, and experimental complexity. Conversely, label-free optical imaging offers substantial technical simplicity but is believed to have little true molecular specificity. Contrary to common belief, here we introduce *super-multiplex* optical imaging without labeling. By systematically studying paired vibrational spectroscopic imaging and mass spectrometry imaging, we discovered a surprisingly strong (more than 0.9) correlation between their latent space representations, supported by both experiments and theory. This insight prompts us to build supervised learning models to successfully predict spatial distribution of 100 molecular species directly from label-free vibrational images across diverse tissue systems. We developed this technology, named *Prediction through Learning with AdvaNced Chemical Kaleidoscope* (PLANCK), and demonstrated it with both infrared-based vibrational imaging of organ-scale tissues and Raman-based vibrational imaging of live tissues. Powered by AI, PLANCK decodes the exquisitely rich but otherwise hidden vibrational information into a surprisingly large number of ( ≥ 100) specific molecular species, providing a cost-effective and scalable solution for basic research and translation, including applications in live imaging.

## Introduction

Optical imaging has transformed modern biology. While recent advances have reached nanometer spatial resolution, microsecond time resolution and organism-level depth penetration, the amount of molecular specific information has remained difficult to expand, limiting the ability to visualize a large set of specific biomolecules. This has become a severe bottleneck in the age of omics. Thus, how to achieve super-multiplex optical imaging of specific molecules has emerged as the next frontier of innovation (Fig. 1).

**Fig. 1.**
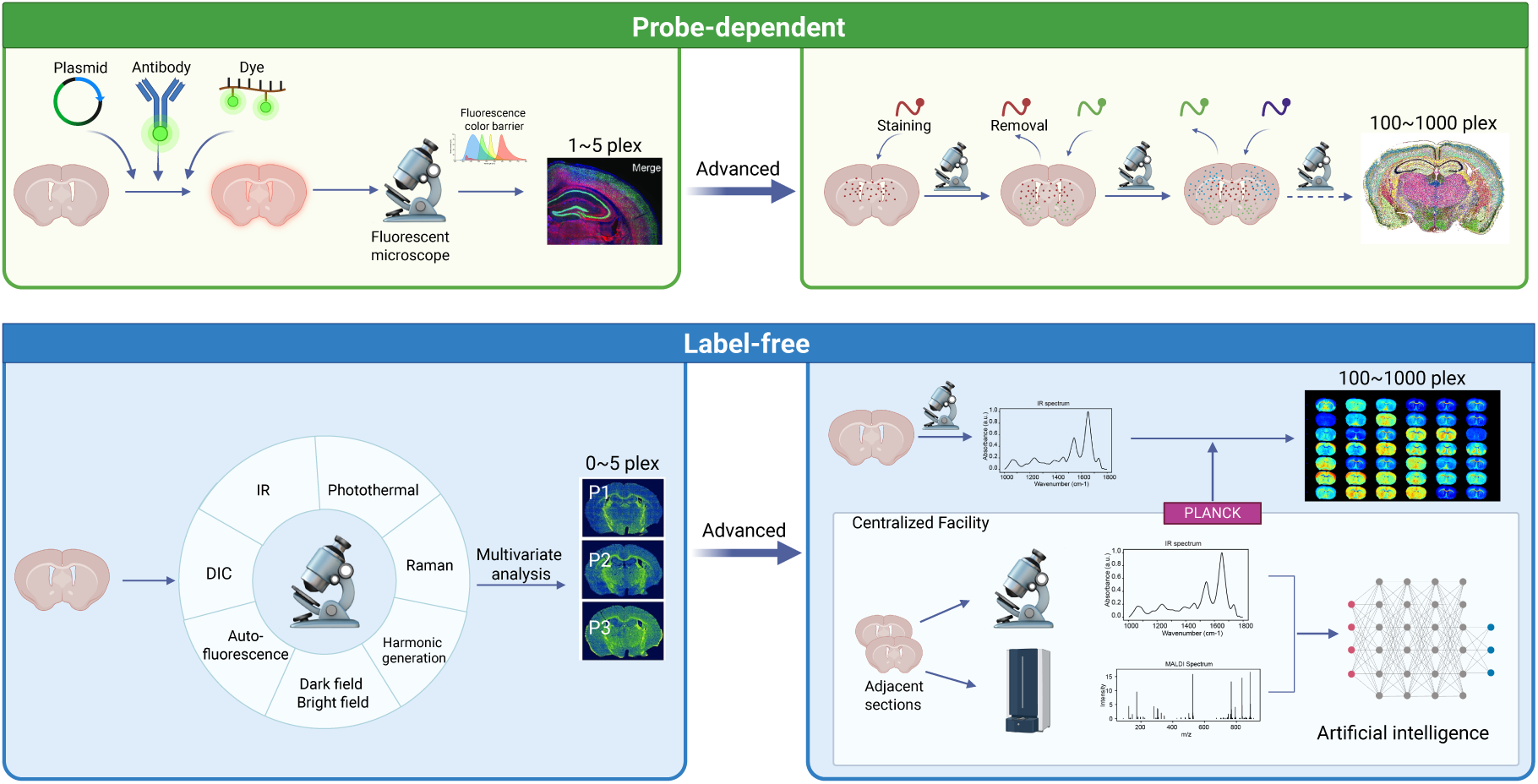
Principle illustration of PLANCK. Summary and comparison of optical imaging techniques in the context of multiplex imaging. **a**, Traditional multi-color optical (fluorescence) imaging through probe labeling, with limitations of 3∼5 colors due to fluorescence’s color barrier. **b**, Sophisticated labeling techniques for multiplex optical imaging (100-1000 plex), which often requires high-cost instrument and reagents and complex sample preparations via probe cycling or barcoding. **c**, Label-free optical imaging with limited molecular information (0-5 plex). **d**, PLANCK: label-free super multiplex optical imaging (100 – 1000 plex), achieved by paired measurements between vibrational imaging and molecular-specific imaging first (MALDI selected here as an example) followed by machine learning. Leveraging the power of AI and cloud technologies with continuing upgrades, the deployment of PLANCK is expected to meet minimum barrier among the wide community.

The molecular specificity has almost always been conferred via labeling prior to imaging. By tagging specific biomolecules with fluorophores (such as fluorescent dyes, proteins and antibodies), researchers can achieve high molecular specificity. However, the broad and featureless fluorescence spectrum limits the number of simultaneous imaging to less than 4∼5 colors (Fig. 1a). To overcome this color barrier, sophisticated techniques such as vibrational probes^1–3^, cyclic labeling^4–6^, combinatorial barcoding^7–9^ have enhanced the multiplexing level to 10-100 molecular species or even greater (Fig. 1b). Despite their remarkable success, these labeling-focused methods often require expensive instrumentation and reagents, complicated sample treatment (often on fixed samples only) or exceedingly long experimental time. In addition, a significant percentage of biomolecules, especially those small metabolites, remain untaggable by current probes^10^.

Contrary to the use of labels, label-free optical imaging excels in its technical simplicity, bypassing all the experimental difficulty, cost and time associated with preparing, introducing and detecting multiple probes in complex biological environment. However, its molecular information is believed to be drastically poorer as a trade-off. For example, the phase contrast and differential interference contrast microscopy register change of refractive index, which has intrinsic low molecular specificity. Auto-fluorescence can simply map out few natural fluorophores. Photothermal and photoacoustic imaging only report highly concentrated class of light-absorbing species. Second harmonic generation is mostly selective to fibrillar collagen. Vibrational imaging such as Raman and infrared (IR) microscopy can provide information of functional chemical groups. However, as these functional groups are shared among a myriad of biomolecules (in 10,000s) inside cells, vibrational imaging usually only identifies gross classes of biomolecules such as total proteins and total lipids instead of specific molecular species^11–13^. Generally speaking, the true molecular–level multiplexing of label-free optical imaging is ranging between 0 and 5 in most circumstances (Fig. 1c).

Herein we report a new technology to achieve super-multiplexed optical imaging without labeling. Vibrational spectroscopy based chemical imaging has found increasing utility in phenotyping cell states and tissue pathology^12,14–21^. These successful applications imply that, while vibrational spectrum of biological samples might seem highly congested and obscure, it may contain a significant amount of information, albeit not readily interpretable. To investigate the hidden information, we performed paired measurements between infrared (IR) microscopy and mass spectrometry imaging, a molecularly informative but non-optical technique^22–25^. Our analysis revealed surprisingly high correlation and similar latent space distributions between these two measurements. This discovery prompts us to develop supervised machine learning models to decode the vibrational spectroscopy that is hardly decipherable otherwise. The trained model can perform spectral decoding and generate 100-plex molecular images cross the organ-scale tissue with high performance (average correlation of 0.86). Moreover, we extend this strategy to Raman imaging: the combination of state-of-the-art stimulated Raman scattering microscopy^26–28^ and specifically designed deep learning model (Hyper-pix2pix) enables super-multiplexed imaging of live samples, *for the first time*. Our technology is termed *<u>P</u>rediction through <u>L</u>earning with <u>A</u>dva<u>n</u>ced <u>C</u>hemical <u>K</u>aleidoscope* (PLANCK), as it taps into the information wealth of chemical imaging. Given the simplicity of label-free imaging, and that the machine learning models could be pre-trained in centralized facility and continuously upgraded in cloud, its deployment is expected to meet minimum barrier among the wide community (Fig. 1d).

### Vibrational imaging captures functional group information but unclear amount of molecular specific information

Vibrational spectroscopic imaging measures functional chemical groups such as C-H, C=O, C-C, P=O and amide. Its success in phenotyping cell states and tissue pathology imply that the spectrum should contain good amount of biologically relevant information^12,14–21^, albeit not readily interpretable. To showcase the captured information, we chose Fourier transform infrared (FTIR) microscopy as a demonstration, as it is a widely used, commercially available technique for mapping large-scale tissues with cellular resolution^29^. Mouse brain tissues were selected as our model system, due to the rich cell types and cell states in the brain. Hyperspectral FTIR dataset was collected in the fingerprint region. By applying K-means clustering on the collected dataset, the result clearly shows meaningful spatial pattern that is associated with the mouse brain anatomy (Supplementary Fig. 1), supporting that the FTIR dataset captures valuable information.

However, how much molecular specific information the spectrum carries is unclear and debatable. Since common functional groups (several 10s in number) are shared among a myriad of biomolecules (in 10,000s) inside cells (Supplementary Fig. 2), there does not exist a simple one-to-one mapping recipe. Indeed, spectral unmixing can only yield a few rough classes of biomolecules such as total proteins, total lipids and total nucleic acids^11–13^. Even when applying advanced algorithms such as multivariate curve resolution-alternating least squares analysis, the molecular specificity is still limited^30^, especially within complex mammalian tissue systems (Fig. 1c). Thus, it remains elusive whether it is possible to decode or translate the information hidden in vibrational spectroscopic imaging to actual molecular information.

### Surprisingly high correlation and similar latent distribution between vibrational imaging and mass spectrometry imaging

We asked whether vibrational imaging contains molecular information that is not directly assignable from individual vibrational bands but may be recoverable through its relationship with a molecularly specific imaging modality. To do so, we designed paired measurements between FTIR microscopy and mass spectrometry imaging, which can provide ground-truth spatial distribution of a large set (often 100∼1000) of specific molecules such as metabolites, lipids, carbohydrates and peptides^22–24^. Among common sample ionization methods, matrix-assisted laser desorption/ionization (MALDI) is one of the most popular methods because of its high sensitivity and spatial resolution (10-100 microns), wide molecular coverage, reliable identification and annotation^31–33^. The workflow of our paired measurements is illustrated in Supplementary Fig. 3. Fresh frozen mouse brain tissues were sectioned in adjacent slices and prepared separately for MALDI and vibrational imaging using each modality’s optimized protocol, to ensure optimal data quality for both modalities. After imaging, both hyperspectral datasets underwent standard data preprocessing, and a subset of MALDI peaks (100 m/z peaks) with clear patterns and biological annotations was selected for downstream analysis. The two image volumes were then registered, yielding aligned multimodal data for subsequent analysis and prediction (Methods).

Because both IR and MALDI imaging datasets are intrinsically multivariate, traditional univariate analysis is not suited. In other words, comparing individual IR bands with individual m/z peaks is insufficient to evaluate their relationship. To address this, we employed canonical correlation analysis (CCA), a classical statistical method that measures the associations of two multivariate datasets^34^, to test whether IR and MALDI share a common low-dimensional latent structure of possessed molecular information. CCA identifies paired linear projections of the IR spectral space and MALDI molecular space, denoted as U and V, whose scores are maximally correlated (Fig. 2a).

**Fig. 2.**
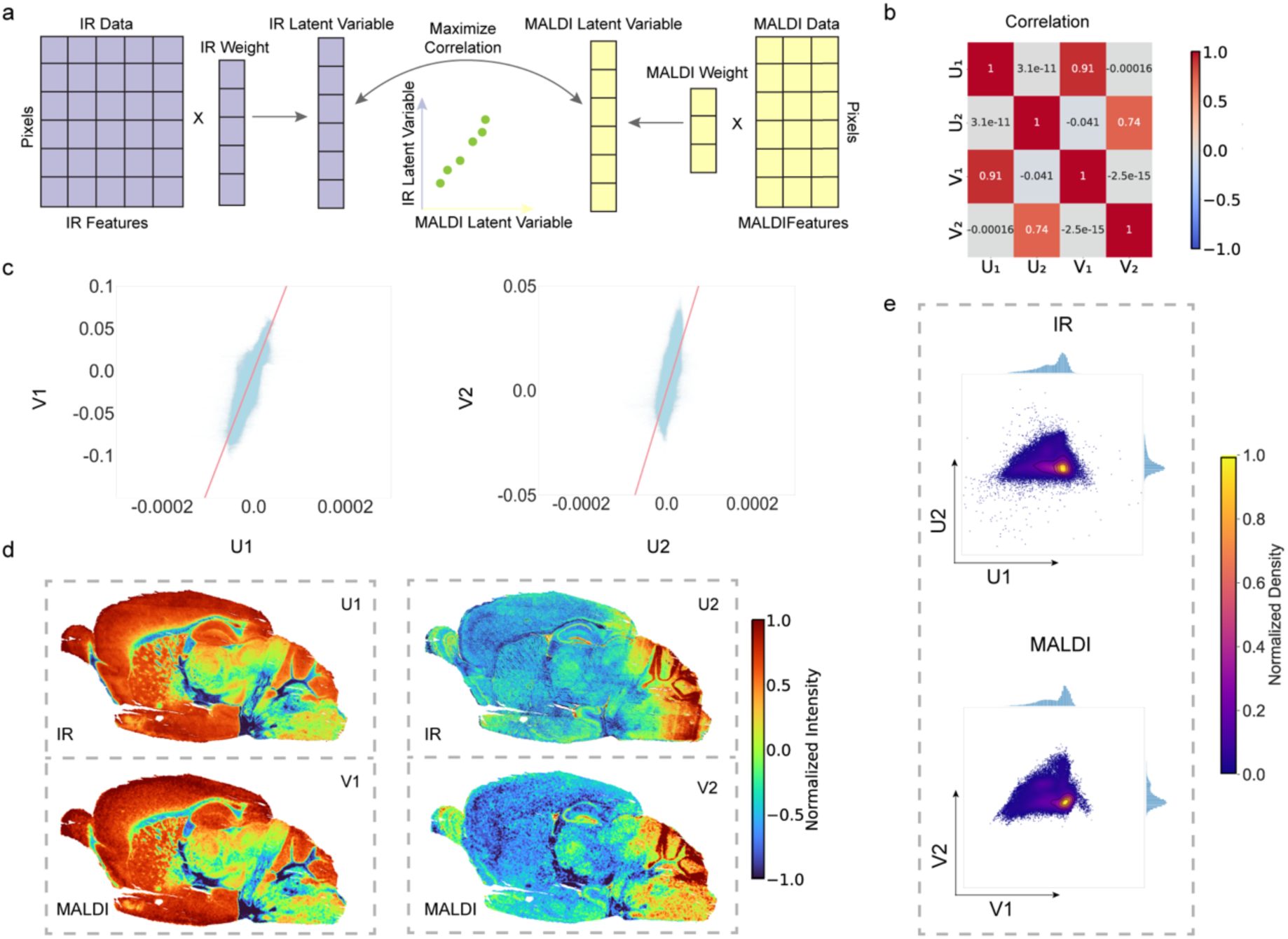
Canonical correlation analysis (CCA) reveals the high correlation and similar latent distribution between FTIR imaging and MALDI imaging. **a**, CCA principle of analyzing the correlation of IR and MALDI datasets. **b**, Correlation heatmap of the top 2 pairs of latent canonical variables of IR (U_1_, U_2_) and MALDI (V_1_, V_2_) in CCA, demonstrating high correlation of canonical variables across the two datasets and the orthogonal relationship of canonical variables within one dataset. **c**, Scatter plots of top 2 latent variables between IR and MALDI (U_1_ vs V_1_, U_2_ vs V_2_), showing nearly linear association. **d**, Real-space visualization of top 2 canonical variables’ scores in both IR and MALDI dataset, showing high similarity. **e**, Low-dimensional data distribution of IR and MALDI when projected to their own latent variables (U_1_, U_2_) and (V_1_, V_2_), respectively, showing highly resembled latent space. Color scale: normalized intensity of signal at latent space.

Using 207 IR spectral features and 100 MALDI m/z channels, CCA revealed strong cross-modal correspondence. The first pair of canonical variates (U_1_ and V_1_), reached a correlation of 0.91, and even the second pair remained highly correlated at 0.74 (Fig. 2b). The matched scatter plots showed near-linear relationships between the corresponding IR and MALDI-derived canonical variates (Fig. 2c). When these scores were mapped back to tissue space for visualization, the FTIR and MALDI canonical images showed highly similar spatial patterns (Fig. 2d), indicating that independent combinations of vibrational bands and molecular ions capture related biochemical variation at tissue-scale.

We further examined whether this correspondence was preserved at the level of latent data organization. Because the canonical variables define low-dimensional representations of each modality, we projected the IR and MALDI data into their respective first two canonical dimensions, (U_1_, U_2_) and (V_1_, V_2_). The resulting distributions were closely matched (Fig. 2e), suggesting that the two modalities not only produce similar spatial patterns, but also preserve similar organization of biochemical/molecular states in latent space. This latent-space correspondence was reproduced in additional paired datasets across tissue sections and tissue types (Supplementary Fig. 4), whereas the high correlations and matched latent distributions disappeared when IR and MALDI datasets were unpaired or randomly simulated (Supplementary Fig. 5). Together, these results show that vibrational imaging and MSI share a very similar latent molecular structure, supporting the feasibility of using molecularly specific MSI measurements to inform and decode label-free vibrational imaging.

### Regression analysis connects IR spectrum to 100-plex molecular species

Having shown that IR and MALDI share a similar latent structure, we next examined whether this relationship could be expressed as an explicit transformation from vibrational spectra to specific molecular channels. This analysis aimed to determine whether a learned spectral-to-molecular transformation can quantitatively connect the two modalities and reveal chemically meaningful relationships between IR features and MALDI m/z channels.

Considering the matrix nature of two datasets, a natural way to formulate this problem is to learn a transformation from the IR spectral matrix to the MALDI molecular matrix. A transformation matrix, T, can then be estimated to map the IR spectral space onto the MALDI molecular space (Fig. 3a). This formulation provides a general mathematical view of molecular decoding, in which regression models can approximate the spectral-to-molecular transformation from paired measurements.

**Fig. 3.**
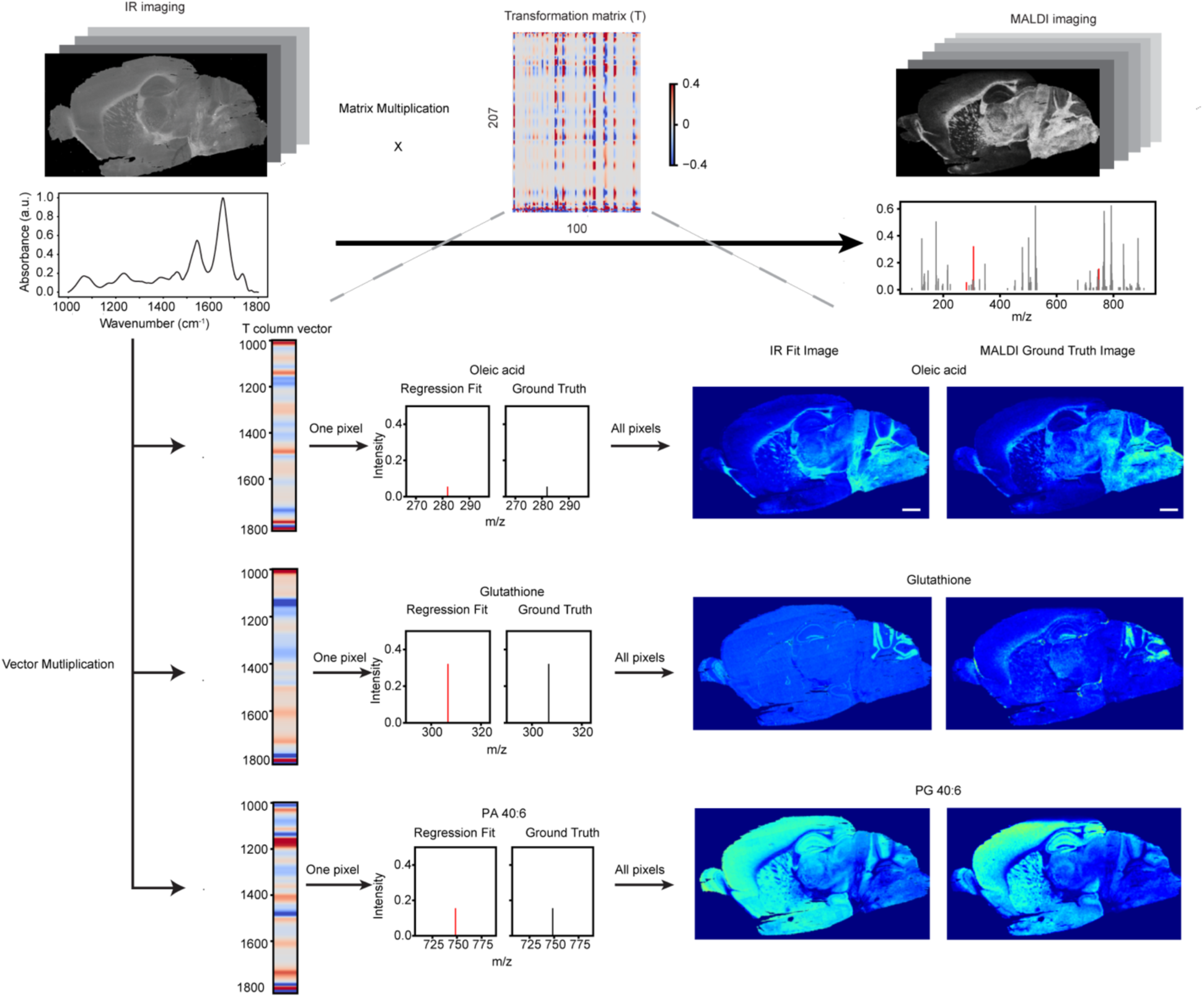
Regression analysis on paired IR and MALDI datasets, connecting two datasets through matrix transformation. IR data can be decoded into MALDI data through a fitted transformation matrix, T, in regression analysis. Three examples (oleic acid, glutathione and PG 40:6) were used to showcase that the dot product between an IR spectrum vector of a single pixel and a column vector in the transformation matrix, T, represents the MALDI intensity signal of a given biomolecule at that pixel. When this calculation is applied to all pixels, it produces the entire fitted image of the given biomolecule, which matches well with the ground truth MALDI image. For each column vector, red color in the middle corresponds to IR peaks with larger weights in transformation of a given biomolecule. Note that while the column vectors’ two extreme ends often appear largely weighted, they hold little biological significance due to their near-zero IR intensity. Scale bar: 1mm. Color scale: signal intensity in transformation matrix.

Using this fitted transformation, the 100 selected MALDI m/z channels were reconstructed from the corresponding IR spectra with high agreement to the measured MALDI maps, achieving an average correlation of 0.9018 across channels (Supplementary Fig. 6). This result indicates that the paired IR–MALDI data contain sufficient shared structure for a learned transformation to connect label-free vibrational spectra with molecularly specific MSI channels.

We next visualized this transformation using representative molecular channels, including oleic acid, glutathione, and PG 40:6 (phosphatidylglycerol), whose corresponding m/z peaks are highlighted in Fig. 3. For each molecular species, the corresponding column of the transformation matrix defines a weight vector that maps IR spectral features to the intensity of that MALDI molecular channel. Multiplying the IR spectrum at each pixel by this weight vector generates a scalar value corresponding to the reconstructed MSI intensity for that molecule (Fig. 3, middle). Repeating this operation across all pixels produces a reconstructed molecular image, which can be directly compared with the measured MALDI image (Fig. 3, right). For these representative molecules, the reconstructed spatial patterns closely resembled their MALDI references, illustrating how the transformation matrix links label-free vibrational spectra to molecularly specific MSI maps.

The transformation matrix also provides insight into the spectral basis of this cross-modal relationship. Each column of the matrix links IR spectral features to one MALDI m/z channel. For example, the oleic acid-associated weights were enriched near 1465 cm^-1^, corresponding to lipid CH_2_ bending vibrations (*δ*CH_2_) in IR spectrum, consistent with its fatty-acid structure. Glutathione showed stronger weights in the 1550–1650 cm^-1^ region, associated with protein amide vibrations and consistent with its peptide backbone. PG 40:6 showed prominent weights around 1150–1170 cm^-1^, a region associated with C–O vibrations in carbohydrates/glycerol. These examples suggest that the fitted FTIR-to-MALDI transformation is not arbitrary, but contains chemically interpretable spectral features that connect vibrational contrast to molecularly specific MSI channels.

Together, this transformation analysis demonstrates that IR spectra and MALDI molecular channels can be quantitatively connected through a chemically interpretable spectral-to-molecular mapping. This provides the mathematical and interpretability basis for PLANCK, while the following section tests whether this relationship can be transferred beyond the fitted paired dataset to predict molecular maps across tissues.

### PLANCK predicts 100-plex molecular species from IR spectrum across tissues

Having established a spectral-to-molecular transformation between IR and MALDI, we next asked whether this relationship generalizes beyond the fitted samples. Specifically, we investigated whether a transformation learned from one set of paired tissues could decode molecular information from unseen IR data acquired from independent tissue sections. To this end, we implemented PLANCK using partial least squares (PLS) regression^35^, a classical multi-output regression method that projects the input IR spectra into latent variables optimized to maximize covariance with the target MALDI molecular channels. This covariance-based formulation is consistent with the strong cross-modal correlations identified by CCA, reinforcing the suitability of PLS for decoding and predicting molecular information from label-free IR imaging.

We collected 6 pairs of mouse whole-brain sagittal sections, 2 pairs of sagittal cerebellum sections, and 2 pairs of whole-brain coronal sections. Each whole-brain sagittal dataset contained approximately 8 million spectra. The data were split into 6 paired sections for training and 4 paired sections for testing, with the test set including 2 whole-brain sagittal sections, 1 sagittal cerebellum section, and 1 whole-brain coronal section. The PLS model was trained exclusively on the training dataset and then applied to the test dataset for evaluation. This design enabled us to evaluate PLANCK across independent tissue sections, anatomical orientations, and brain tissue types.

PLS-based PLANCK effectively learns to predict 100-plex molecular channels across different brain tissue types (Fig. 4b-g). For testing data of whole-brain sagittal sections, five m/z peaks with distinct spatial patterns were selected for demonstration (Fig. 4b). The PLANCK-generated molecular maps reproduced major anatomical and regional patterns observed in the corresponding MALDI reference images. At the spectral level, the average PLANCK-estimated molecular profile showed strong agreement with the MALDI spectrum (Fig. 4c), with only minor residual deviations (residual plot in Supplementary Fig. 7). For quantitative evaluation, we selected peak signal-to-noise ratio (PSNR), structural similarity index (SSIM), and Pearson correlation coefficient (PCC)^36^. Boxplots summarizing 100 m/z peaks are shown in Fig. 4d, with averages of 24.03 dB (PSNR), 0.8995 (SSIM), and 0.8610 (PCC) in whole-brain sagittal test datasets. For context, studies predicting spatial-omics based on H&E images usually yield PCC in the range of 0.2-0.6^37–39^, underscoring the advantage of vibrational imaging as a molecularly informative predictive modality.

**Fig. 4.**
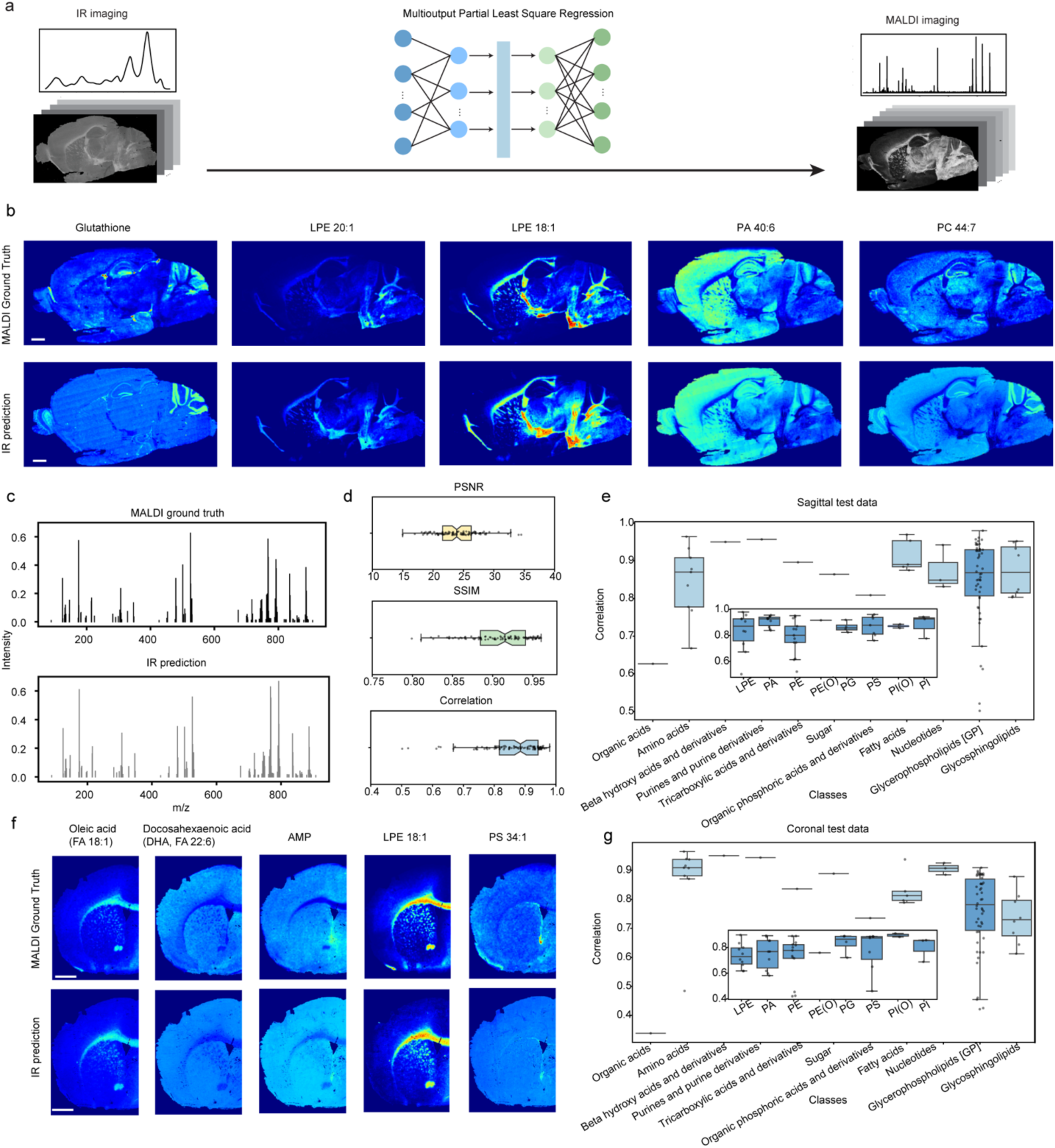
PLANCK: partial least square (PLS) regression decodes IR imaging into MALDI imaging with 100-plex specific molecular information in mouse brain systems. a, Illustration of the decoding mechanism from IR to MALDI via PLS. **b**, Demonstration of 5 predicted biomolecular images predicted from IR, showing highly similar patterns to the MALDI ground truth. Scale bar: 1mm. **c**, Average mass spectrum from IR prediction and MALDI ground truth, showing strong alignment. **d**, Quantitative evaluations of IR prediction performance using PSNR, SSIM and correlation metrics, with average of 24.03 dB (PSNR), 0.8995 (SSIM), and 0.8610 (PCC). The boxplots summarize the prediction performance across 100 m/z peaks in 2 whole-brain sagittal sections in the testing dataset. **e**, Correlation performance of 11 classified biomolecular categories in whole-brain sagittal sections. **f**, Demonstration of 5 predicted biomolecular images in mouse whole-brain coronal section, indicating the generality of PLANCK. Scale bar: 1mm. **g**, Correlation performance of 11 classified biomolecular categories in whole-brain coronal sections.

The prediction performance of PLS-based PLANCK slightly varied across different molecular species, indicating that PLANCK does not infer all MALDI channels equally. Among specifically MALDI annotated biomolecular groups (Fig. 4e), predictions were most accurate for certain lipids (beta hydroxy acids, fatty acids, tricarboxylic acids, phosphosphingolipids), nucleotides and purines, which have pronounced vibrational peaks in IR spectra. Other lipids (glycerophospholipids/GP, glycosphingolipids) and amino acids exhibited prediction performance with larger variations. The insert figure includes the prediction performance across 8 GP subgroups, demonstrating PLANCK’s predictive capabilities in more refined molecular categories. For other metabolites such as organic acids and organic phosphoric acids, their prediction performances are relatively lower. These differences suggest that PLANCK estimates molecular channels according to the strength of their spectral, spatial, and biological associations with the IR measurements.

To assess method generalizability, we evaluated its performance on held-out mouse coronal sections (Fig. 4f-g, Supplementary Fig. 8) and sagittal cerebellum sections (Supplementary Fig. 9). The PLS model maintained strong predictive performance in these independent samples, such as the mouse-brain coronal section. This was supported by consistent predicted image and spectral profiles, as well as quantitative evaluations (Fig. 4f-g). Because sagittal whole-brain, sagittal cerebellum, and coronal brain sections contain overlapping but partially distinct anatomical structures, cell populations, and molecular compositions, these results support PLANCK’s ability to generalize beyond individual paired sections. Importantly, this validation establishes cross-section, cross-orientation, and cross-region generalization within mouse brain tissues, while broader cross-organ generalization and direct biochemical validation remain important directions for future investigation.

### Super-multiplex live imaging through Raman microscopy and deep learning in PLANCK

We then extend the generality of PLANCK from IR imaging to Raman imaging, another technological feat. Compared to IR microscopy, Raman imaging offers subcellular spatial resolution and compatibility with live imaging. Hence it would be highly desirable to perform super-multiplex live imaging through Raman modality via PLANCK. However, the standard spontaneous Raman imaging is too time-consuming for large-scale tissue imaging. Harnessing quantum amplification, stimulated Raman scattering (SRS) is able to provide orders of magnitude higher sensitivity and imaging speed^26–28,40^. The standard narrowband SRS microscopy only addresses a single vibrational mode, which sacrifices spectral information. To address these challenges and balance the trade-offs, we employed multi-color SRS imaging, focusing on the most pronounced and also spectrally informative C–H vibrational peaks (Supplementary Fig. 10) as a demonstration of PLANCK in Raman microscopy modality (Fig. 5a).

**Fig. 5.**
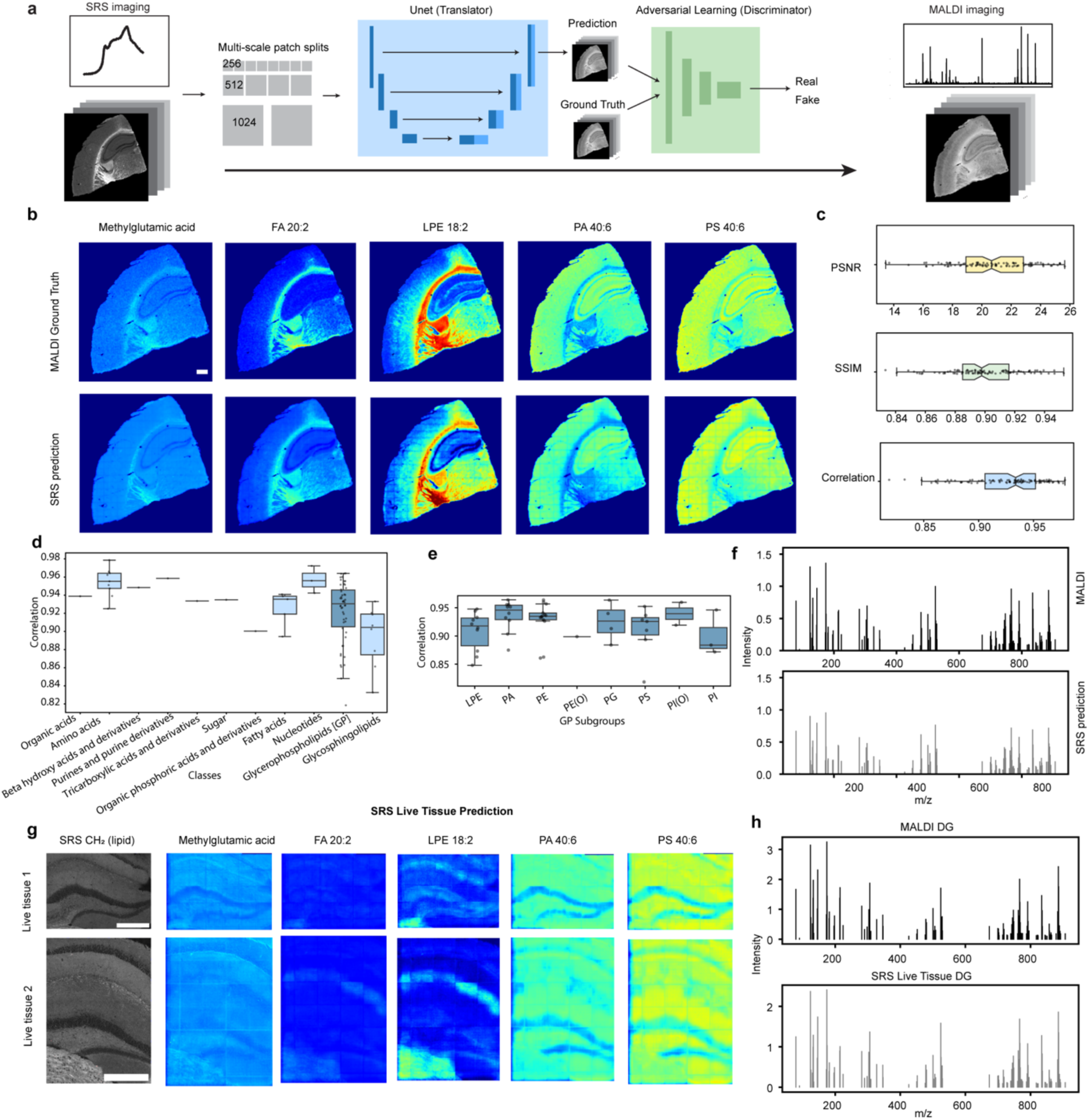
PLANCK: deep learning decodes SRS imaging into MALDI imaging with 100-plex specific molecular information in fixed and live samples. a, Illustration of the decoding mechanism from SRS to MALDI via a designed deep learning model (Hyper-pix2pix). **b**, Demonstration of 5 biomolecular images predicted from SRS, showing highly similar patterns to the MALDI ground truth. Scale bar: 500 microns**. c**, Quantitative evaluations of SRS prediction performance using PSNR, SSIM and correlation metrics, with average of 20.69 dB (PSNR), 0.8994 (SSIM), and 0.9259 (PCC). The boxplots summarize the prediction performance across 100 m/z peaks in all quarter coronal sections in the testing dataset. **d**, Correlation performance of 11 classified biomolecular categories in mouse quarter coronal sections. **e**, Correlation performance of 8 GP subgroups in mouse quarter coronal sections. **f**, Average mass spectrum from SRS prediction and MALDI ground truth, showing strong alignment in spectrum profiles. **g**, Demonstration of predicted biomolecular images in 2 live mouse quarter coronal samples, indicating the ability of PLANCK for super-multiplex live imaging. Left are SRS lipid images and right are predicted 5 images from the same biomolecules in **b**. Scale bar: 500 microns. **h**, Average mass spectrum in the dentate gyrus (DG) region from SRS prediction in live sample 1 and MALDI in another fixed sample.

We found PLS to be less effective in decoding SRS images into multiplex molecular images due to the reduced spectral features (Supplementary Fig. 11). To address this, we designed a deep learning model that leverages the high-resolution spatial features as well as spectral features of SRS images for more accurate translation (Fig. 5a). Our model builds upon the pix2pix architecture^41^, an emerged conditional generative adversarial network (cGAN) model for image translation, but incorporates several key innovations specifically designed for hyperspectral SRS image translation. We refer to this enhanced model as Hyper-pix2pix. The main modifications include (1) the use of group normalization to better handle spatial and spectral features; (2) loss function modification with addition of a spectrum loss to reconstruct spectral profiles; and (3) an ensemble learning strategy that captures spatial features across multiple spatial scales by dividing large-scale tissue sections into multi-scale patches (Methods). Additionally, unlike typical GAN setups, we omit random noise in the inputs, allowing the model to generate discriminative outputs.

We used mouse quarter-coronal brain tissue sections for demonstration. We collected paired datasets from 7 tissue sections, using 5 for training and validation (85% training, 15% validation) and 2 for testing. The training set contains 1465, 386 and 106 images for patch size at 256, 512 and 1024, respectively, with additional data augmentation. The Hyper-pix2pix model was trained only on the training dataset and then applied to the test sections without refitting or using MALDI information from the test samples.

Under this setting, Hyper-pix2pix generated MALDI-referenced molecular maps from SRS images, with performance summarized in Fig. 5b–f. Similar to the IR results, five biomolecules with distinct spatial patterns were selected for visualization (Fig. 5b), where the predicted molecular images closely resembled the MALDI ground truth. Quantitative evaluations (Fig. 5c) showed an average PSNR of 20.69 dB, SSIM of 0.8994, and a correlation of 0.9259 across 100 molecular species in test data. Spectrally, the averaged mass spectra estimated from SRS showed strong agreement with the MALDI spectra (Fig. 5f) although the predicted spectra exhibited slightly lower intensity overall (residual plot, Supplementary Fig. 12)

Across annotated biomolecular groups (Fig. 5d), the performance trend was broadly similar to that observed in the IR-based analysis, with stronger performance for biomolecules related to carbohydrates, nucleic acids, and most lipid classes. Notably, estimation of some small metabolites, including organic and tricarboxylic acids, improved in this setting. Performance across 8 glycerophospholipid subgroups is summarized in Fig. 5e. Direct performance comparison between Hyper-pix2pix, pix2pix, U-Net^42^, UwU-Net^43^ and PLS (Supplementary Table 1) further demonstrated the advantages of our tailored Hyper-pix2pix model for hyperspectral image translation in PLANCK.

By incorporating Raman microscopy modality, PLANCK enables super-multiplex optical imaging in live tissue samples—an advantage unachievable with mass spectrometry imaging (or other omics technology so far) due to its destructive nature. To demonstrate this possibility, we collected SRS images from live mouse quarter-coronal sections (Fig. 5g–h), capturing only subregions to maintain sample viability during imaging. As expected, Raman spectra are identical in this region between fixed and live tissues (Supplementary Fig. 13), making training on fixed tissues viable to predict live tissues. Two live samples are shown in Fig. 5g, with the leftmost images displaying original SRS lipid images and the right five images presenting predicted molecular distributions. The predicted images preserve the anatomical structures seen in the SRS images. Additionally, they correspond to the same biomolecules shown in Fig. 5b, maintaining similar contrasts and patterns as observed in fixed samples. Spectrally, the average 100-peaks-spanning spectrum predicted from the dentate gyrus (DG) region of the live samples closely resembles that of the MALDI sample which has to be fixed, demonstrating strong spectral consistency (Fig. 5h). Overall, these results highlight PLANCK’s ability to generate biologically meaningful molecular images in live samples.

### Theoretical rationale underlying PLANCK

To understand the theoretical link between vibrational imaging and mass spectrometry imaging (MSI), we propose a theoretical framework to elucidate their connection. It is well established that both modalities exhibit a linear concentration dependence in analyte response. In an ideal scenario, the imaging data can be mathematically expressed as the multiplication of two matrices: a pixel-independent concentration matrix of biological analytes and a modality-specific response matrix (Equations 1–2). Here, we denote the matrix form of vibrational imaging as **E**, considering it measures the vibrational energy of chemical bonds. **E** is in dimension of *N* × *n*_e_, where N is the total number of pixels and *n*_e_ is the dimension of vibrational spectrum. Similarly, MSI data is represented as **M**, with dimensions of *N* × *n_m_*, where *n_m_* is the dimension of mass spectrum. The concentration matrices **C** and ***C***′ represent biological analyte concentrations detectable in vibrational imaging (with a total number of *n*_c_) and MSI (with a total number of *n*_c_′), respectively, while ***R_e_*** and ***R_m_*** represent the corresponding modality-specific response matrices. A detailed derivation of Equations 1–2 is provided in Methods.

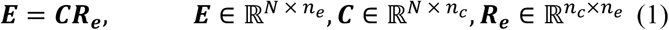

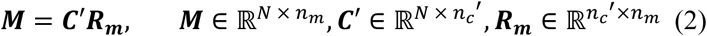

A central question is whether the two concentration matrices ***C*** and ***C***′ are identical in two imaging modalities. If they were, **E** and **M** would be linearly connected by the concentration matrix. We reason that in biological tissues (e.g. brain), they are unlikely to be completely identical or independent, because each modality is sensitive to partially overlapping but distinct sets of molecule species. We therefore decompose the concentration space into shared and modality-specific components. Specifically, ***C_o_*** denotes molecular components that contribute to both vibrational imaging and MSI; ***C_e_*** denotes components that contribute only to vibrational imaging; and ***C_m_*** denotes components that contribute only to MSI. For example, subsets of lipids, nucleotides, and carbohydrates may contribute to both modalities (supported by Fig. 4-5), whereas proteins respond only in vibrational imaging. Conversely, some low-abundance metabolites or ions may be only detected by MSI. Under this decomposition, Equations 1–2 can be written as matrix blocks (equation 3-4), where ***R_e_***_,***o***_ and ***R_m_***_,***o***_ represent the corresponding response of overlapping concentration in **E** and **M**, respectively, and ***R_e_***_,***e***_ and ***R_m_***_,***m***_ represent the response of non-overlapping concentrations.

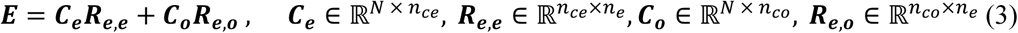

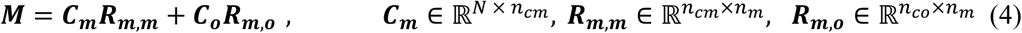

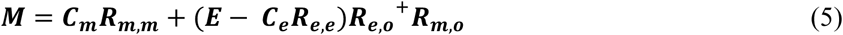

Applying Moore-Penrose pseudoinverse^44^ (denoted as ^+^) provides a least-squares approximate solution to express the overlapping concentration ***C_o_*** in equation 3. When plugging this expression into equation 4, equation 5 can be derived, indicating that **M** can be partially expressed by **E** through matrix transformation (the second term). This illustrates that the MSI measurement can be decomposed into two parts: a component associated with molecular information shared with vibrational imaging, and a remaining MSI-specific component. Thus, the theory does not require that vibrational imaging fully replaces MSI or uniquely determines every MSI channel. Instead, it suggests that MSI-referenced molecular channels are estimable from vibrational imaging when the relevant molecular components either share direct spectral information or are statistically coupled to vibrationally detectable tissue features.

The remaining uncertainty lies in the modality-enriched terms ***C_e_R_e_***_,***e***_ and ***C_m_R_m_***_,***m***_. We reason that ***C_e_*, *C_o_*, *C_m_*** are not completely independent, especially in biological tissue systems. For instance, the large protein concentration in ***C_e_*** may correlate to the amino acid concentration in ***C_o_***; the metabolite concentration in ***C_m_***, such as choline, can be closely related to lipids concentration in ***C_o_***, as choline plays an important role in lipid transport and cell membrane structure. If these dependencies primarily follow linear combinations, **M** can be shown to be largely expressed by **E** through matrix transformation (Methods). This aligns with the strong correlation between two datasets via CCA, as well as strong prediction result achieved by linear regression models such as PLS. If these dependencies are nonlinear, **M** can still be mathematically expressed by **E**, though in a more complexed format (see Methods). We suspect that in this case, the nonlinear operations in deep learning matters to infer **M** from **E**.

The biological intuition behind our theory is that complex biological systems normally operate in relatively low-dimensional spaces^45,46^. Due to the inherent structure of cellular systems, genes are co-regulated and cellular features are interrelated^46^. Thus, despite the vast number of possible cell states in biological systems, many of them are connected and correlated. In fact, it has been suggested that only a countable number of cell states govern the possible high-dimensional geometry of physical measurements^46,47^. Hence it is conceivable that vibrational imaging data and MSI imaging data are fundamentally linked by a finite number of cell states. Moreover, such associations can extend to other imaging modalities within the PLANCK framework.

## Conclusion and Discussion

How to expand the multiplexing level has become a frontier of optical imaging. While the notion of labeling has accomplished tremendously in the past decades, its constraint is not to be ignored. The experimental expertise, cost and time that is required for preparing, introducing and detecting multiple probes in biological environment is not a trivial factor, which only goes up steeply as the demand of multiplexing increase further^1–9^. In this sense, PLANCK takes an opposite route and circumvents these limitations altogether. The choice of molecular specific information is provided by the learning modality in PLANCK as opposed to the type of probes (Fig. 1).

This inference-based paradigm of PLANCK is analogous to the prediction paradigm of AlphaFold^48,49^. Though appearing to be less accurate and trustworthy compared to direct physical measurements in the initial stage, it can grow into a powerful tool with high prediction accuracy with increasing data collection and iterative model improvements. Considering that AI-based computational models could be pre-trained in centralized facilities or in cloud service, its deployment is expected to meet minimum barrier and open up new space of applications such as those in low-resource settings.

PLANCK is built upon the wealth of chemical information of vibrational-based chemical imaging. This type of information has proved pivotal in the emerging applications in single cell metabolism, cell phenotyping, tissue pathology^12,14–21^. While these prior applications have used the chemical information either directly or in an unsupervised learning (Supplementary Fig. 1), PLANCK employs a supervised learning to decode the precious molecular-specific information that is otherwise hidden. The somewhat surprising discovery is that the biological vibrational spectrum can be accurately decoded into concentration information for a remarkably large number of (≥100) specific molecular species, supported by both machine learning results and the analytical theory.

PLANCK can be rather versatile and general, although we developed it through FTIR and SRS microscopy here. Vibrational imaging has grown into a large family of technology with varying capabilities such as video rate live imaging and super-resolution imaging^50–53^. All these capabilities can be potentially equipped with PLANCK.

Importantly, PLANCK is not intended to replace direct molecular measurements by MSI. Rather, MSI serves as a molecularly specific reference modality that enables PLANCK to learn associations between vibrational signatures and MSI-referenced molecular channels. Once trained and validated, PLANCK can generate molecularly informative maps from label-free vibrational images, offering practical advantages in settings where MSI is limited by destructive measurement, matrix coating, ionization, vacuum instrumentation, cost, or incompatibility with live imaging. For example, the estimated cost of FTIR imaging is about 50-100 times lower than that of MADI imaging for the same sample (Supplementary Table 2). Moreover, next-generation IR imaging can be orders of magnitude faster than FTIR^54–56^. Furthermore, SRS is widely used for live tissues, organisms and even humans^50,57^. These features support PLANCK as a complementary strategy for scalable, label-free, molecularly informed optical imaging.

This idea of pairing vibrational imaging with another molecular specific technology via supervised learning is certainly not limited to mass spectrometry imaging only. We expect that PLANCK shall be generalizable to other domains such as spatial transcriptomics and proteomics. In fact, recent studies have suggested connection between Raman and IR spectroscopy and RNA expression^58,59^, although not at imaging setting. These future explorations have the prospect to further mine the information wealth contained in biological vibrational spectroscopy, enlarging the molecular diversity.

## Acknowledgements

We thank Dr. Sami Sauma and Dr. Patrizia Casaccia from the Neuroscience Initiative CUNY-ASRC for providing mouse brain samples. We thank Peiyang Liu, Jiazhang Chen and Rinat Abzalimov for providing technical support on MALDI MSI. W.M. acknowledges support from the National Institute of Health (R35 GM149256) and Chan Zuckerberg Initiative (Dynamic Imaging 2023-321166). Y.H. acknowledges support through PSC-CUNY Faculty Research Award jointly funded by the Professional Staff Congress and The City University of New York, and through NIH shared instrument grant S10OD036268.

## Author Contribution

Xinwen Liu led the theoretical and computational development of PLANCK, including the mathematical theory connecting vibrational imaging and MSI, and the full computational pipeline. Specifically, Xinwen Liu developed the IR, MALDI, and SRS data preprocessing and analysis workflows; established the algorithmic pipeline for multimodal image dataset registration; selected and implemented CCA to investigate relationships across multimodal imaging datasets; designed and performed statistical analysis, regression analysis, PLS-based ML modeling, and Hyper-pix2pix deep learning modeling. Xuemeng Li collected the FTIR imaging data and contributed to CCA analysis and partial dataset registration between SRS and MALDI data. Lele Xu collected the MALDI imaging dataset and performed MALDI metabolite annotation. Mian Wei collected the SRS imaging data and spontaneous Raman data, including both fixed and live samples. A.N. performed froze tissue section and collection of partial MALDI imaging data. Y.H. provided expertise and supervision for the MALDI imaging, including guidance on MALDI sample preparation, tissue sectioning, data acquisition, spectrum annotation, and the incorporation in PLANCK. Xinwen Liu and W.M. conceived the central idea of PLANCK framework, and wrote the manuscript with input from all authors.

## Author Notes

This work was first submitted for journal consideration in Oct 01, 2025.

## Conflicts of Interest

Columbia University (Xinwen Liu and W.M.) has filed a provisional patent application based on this work.

## Methods

### Materials

High purity grade N(1-Naphthyl) Ethylenediamine Dihydrochloride (NEDC, Cat#222488), Phosphorus red (Cat#343242), and carboxymethyl cellulose (CMC, Mw ∼ 90,000; Cat#419273) were purchased from Millipore Sigma-Aldrich (USA). Optima UHPLC/MS-grade Methanol, Isopropanol and water and Poly-D-lysine (PDL, Cat#MP215017580) were purchased from Fisher Scientific (USA). Conductive indium tin oxide (ITO)-coated glass slides (Cat# 8237001) were purchased from Bruker Daltonics (Germany). Methanol (34860-100ML-R) were purchased from Sigma-Aldrich. CaF2 IR Grade (76mm x 26mm x 1mm) were purchased from Crystran.

### Animals

Mice were housed in groups of 4 on a 12-hr light/dark cycle and an ambient temperature of 22 ± 3°C, with food and water available ad libitum. Wildtype male C57BL/6J mice ages 5 weeks were provided by Dr. Patrizia Casaccia laboratory. Animal protocols were approved by IACUC at CUNY Advanced Science Research Center CMU and performed according to NIH guidelines. For live Raman imaging experiments, wildtype mice (C57BL/6, ∼1 month old, Jackson Lab) housed at Columbia University were used. The animal experimental protocol (AC-AABN0554) was approved by the Institutional Animal Care and Use Committee at Columbia University.

### Sample Preparation

For MALDI imaging experiments, animals were sacrificed with cervical displacement and the brains were harvested and frozen for 5 minutes on an aluminum boat floating over liquid nitrogen as previously described [1]. The tissues were stored at-80°C until cryosectioning. Prior to sectioning, CaF₂ slides were coated with 0.1mg/ml poly-D-lysine (PDL) for 2 hours to enhance tissue adhesion. After coating, slides were rinsed twice with distilled water and air-dried at room temperature.

For PLANCK paired measurement, the frozen mouse brains were cryosectioned at 14-μm-thick paired serial sections onto CaF_2_ and ITO slides respectively for MALDI and IR imaging, or three adjacent sections mounted onto CaF_2_, ITO and regular glass slides for IR, MALDI and SRS imaging. This ensures each imaging modality to be operated in its optimal protocol without compromise. Cryosectioning was carried out using CryoStar NX50 (Thermo Scientific, USA) with temperature set at −15 °C for both specimen head and the chamber. Paired adjacent brain sections were gently thaw-mounted onto precooled slides.

Total 7 paired sections of coronal hippocampus, 1 paired section of coronal whole brain, 2 paired sections of sagittal cerebellum and 6 paired of sagittal whole brains were collected for training dataset. The planes of the brain sections collected were determined using the Allen Brain Atlas, corresponds proximately to plate 75/132 and 45/132 of the coronal atlas, and 14/21 of the sagittal atlas.

Mounted cryosections on ITO slides were desiccated in vacuum for 45 min at room temperature, followed by matrix deposition using HTX M5 sprayer (HTX LLC., USA) [1]. 10 mg/ml NEDC [2] in isopropanol/water (70/30, v/v) was deposited at a flow rate of 0.05 ml/min and a nozzle temperature of 80 °C for 30 cycles with no drying between each cycle. A spray velocity of 1300 mm/min, track spacing of 2 mm, N_2_ gas pressure of 10 psi and flow rate of 3 L/min, and nozzle height of 40 mm were used.

#### Fixed mouse brain slice preparation

Fresh frozen mouse brain slices obtained above were fixed with 4% paraformaldehyde in PBS for 15 min at room temperature. For FTIR imaging, fixed slices were washed with dd-H_2_O 5 times for 5 min each time, followed by air-dry overnight; for SRS imaging, fixed slices were washed with PBS 3 times for 5 min each time. The slice was then covered with a coverslip and sealed with nail polish.

#### Live mouse brain slice preparation

A slicing solution was prepared by dissolving 220 mM sucrose, 2.5 mM KCl, 1.23mM NaH_2_PO_4_, 26 mM NaHCO_3_, 1 mM CaCl_2_, 6 mM MgCl_2_, and 2.5 mM glucose in MilliQ water and adjusting the pH to 7.3 with NaOH.

An artificial cerebrospinal fluid (aCSF) solution was prepared by dissolving 124 mM NaCl, 3 mM KCl, 2 mM CaCl_2_, 2 mM MgCl_2_, 1.23 mM NaH_2_PO_4_, 26 mM NaHCO_3_, 3 mM glucose in MilliQ water and adjusting the pH to 7.4 with NaOH.

Wildtype mice (C57BL/6, ∼1 month old, Jackson Lab) were fully anesthetized using isoflurane and then sacrificed with cervical displacement. The whole brain was extracted and immersed in the oxygenated (5% CO_2_ and 95% O_2_) slicing solution pre-cooled at 4°C. After ∼10 min incubation, the mouse brain was cut into a small tissue block with a mouse brain matrix. The tissue block was then sectioned into 200 μm thick coronal slices in the oxygenated slicing solution with a Vibratome. The slices were transferred to an oxygenated aCSF solution at 37°C. A live brain slice sample was prepared by sandwiching a slice in the aCSF solution between a glass slide and a coverslip with two 120 μm thick spacers.

### Imaging and spectroscopy

#### MALDI imaging acquisition

MALDI mass spectra were acquired in negative ion mode by MALDI time-of-light (TOF) mass spectrometer Autoflex Speed (Bruker Daltonics, Germany). MS spectra were calibrated using red phosphorus as the standard for all experiments [1]. The laser spot diameters for coronal and sagittal MALDI runs were focused to “Medium” modulated beam profile for both 30 μm and 50 μm raster width. The imaging data for each array position were summed up by 500 shots at a laser repetition rate of 500 Hz. Imaging data were acquired and processed using FlexImaging v3.0 (Bruker Daltonics), which were further export to.mzML format for downstream computational analysis.

#### FTIR imaging acquisition

Agilent Cary 620 Imaging FTIR equipped with an Agilent 670-IR spectrometer and 128 × 128-pixels FPA HgCdTe (mercury cadmium telluride, MCT) detector was used in the transmission mode. A background spectrum was collected on a clean CaF_2_ substrate using 256 scans at 8 cm^−1^ spectral resolution. Sample spectra were recorded using 16 scans for tissues at 8 cm^−1^ spectral resolution. A ×25 IR objective (pixel size, 3.3 μm, 0.81 numerical aperture (NA)) were used for FTIR imaging. The fingerprint region (1000 – 1800 cm^-1^) with 207 spectrum variables were used for data analysis and prediction in PLANCK.

#### SRS imaging acquisition

SRS imaging was performed on an inverted laser-scanning microscope (Olympus FV1200) using a ×25 water immersion objective lens (Olympus XLPlan N, 1.05-NA, MP, working distance = 2 mm). Two synchronized 2-ps lasers (called pump and Stokes beams) with an 80-MHz repetition rate were provided by a picoEmerald S system from Applied Physics & Electronic. The pump beam is tunable from 720–960 nm. The Stokes beam was fixed at 1031.2 nm. The intensity of the Stokes beam was modulated sinusoidally by a built-in EOM at 20 MHz with a modulation depth of more than 90%. Spatially and temporally overlapped pump and Stokes beams were coupled into the laser-scanning microscope. After passing through the specimens, forward-going pump and Stokes beams were collected with an infrared-coated oil condenser (1.4-NA, Olympus). Stokes beams were completely filtered with two high optical density bandpass filters (890/220 CARS, Chroma Technology), and transmitted pump beams were detected by a large-area (10 mm × 10 mm) Si photodiode (FDS1010, Thorlabs). The output current of the photodiode was then sent to a fast lock-in amplifier (HF2LI, Zurich Instruments) for signal demodulation. The laser power was set as P_pump_ = 14 mW and P_Stokes_ = 80 mW. SRS images were acquired with a pixel dwell time of 4 μs. The time constant of the lock-in amplifier was set as 4 μs. The pixel size was set as 1 μm. The wavelengths of the pump beam were set as 790.5 nm (2940 cm^-1^, symmetric CH_3_ stretching), 796.5 nm (2845 cm^-1^, symmetric CH_2_ stretching), 788.5 nm (2973 cm^-1^, asymmetric CH_3_ stretching), and 794.3 nm (2880 cm^-1^, asymmetric CH_2_ stretching). The Multi-Area Time Lapse (MATL) function was used to acquire images of whole tissue slices.

Flat-field correction was used to balance the uneven laser illumination in each field-of-view. Bovine serum albumin (BSA) was dissolved in PBS to a concentration of 20%. A BSA solution sample was made by sandwiching the solution between a glass slide and a coverslip with a 120 μm thick spacer. A SRS image of the solution sample was acquired at 2940 cm^-1^ with the same parameters as images of mouse brain slices. Each image tile of mouse brain slices was divided by the SRS image of the BSA solution to correct the uneven illumination. The resultant tiles were then stitched into an entire image in ImageJ.

#### Spontaneous Raman spectroscopy

Spontaneous Raman spectroscopy was performed using an upright confocal Raman microscope (Xplora, HORIBA Jobin Yvon). Cell samples were illuminated by 532 nm laser (80 mW on sample) through a 50× objective (air, NA 0.75, MPlan N, Olympus). Specific brain regions were located with a bright-field camera. Raman spectra were acquired with an acquisition time of 5s and 10× accumulation. The grating was set as 1800 gr/mm. Both the slit size and the hole size was set as 100 μm.

### Data preprocessing

#### FTIR imaging data

Data preprocessing was performed using both the commercial software Cytospec and home-built MATLAB scripts with the following steps: (1) PCA noise reduction to denoise and reconstruct the spectra; (2) quality test to remove pixels with a low signal-to-noise ratio in the fingerprint region and the cell-silent region; (3) rubber-band baseline correction for spectral correction; (4) min-max normalization.

#### MALDI imaging data

For each dataset, pyimzml package is used to read and load exported MALDI data in.mzML format into python, followed by binning, baseline correction and root mean square (RMS) normalization. Across datasets, data were manually aligned with similar dataset of mouse brain acquired in timsTOF fleX (Bruker Daltonics, Germany) to ensure mass accuracy (10 ppm tolerance) and iteratively refined to minimize baseline offsets and mass shift artifacts.

For metabolite annotation, multiple approaches were employed to achieve comprehensive annotation. First, the calibrated m/z list was analyzed using Metaboscape (Bruker Daltonics, Germany), based on its exact mass (with an expected mass accuracy of < 5 ppm) and isotope patterns. Putative metabolite identities were assigned by matching their m/z against multiple databases, including The Human Metabolome Database (HMDB), METLIN, LIPID MAPS. Second, the m/z list was submitted to the Metabolomics Workbench for in silico analysis. The resulting candidate matches were evaluated based on delta values, which represent the difference between the observed and theoretical m/z. Third, the m/z list were manually matched with a custom-made metabolite library derived from mouse brain metabolomics data using LC-MS generated in the lab and previously published [2]. In total, 100 m/z peaks with distinctive distribution pattern were selected, among which 82 were annotated across all datasets in this study (Supplementary Table 3). These annotations were further supported by cross-referenced with the metabolite distribution pattern from the Mouse Aging Atlas (mouse.atlas.metabolomics.us) [2]. Additionally, a subset of annotations was further validated by cross referencing with previously published MALDI MSI studies [3-5, 8, 9].

After 100 m/z selection, dataset is log transformed and batch corrected using pyComBat [6] to reduce the batch effects. For data analysis and pls prediction, MALDI data is converted back to the original scale to explore linear connection between FTIR and MALDI data. For deep learning modeling, the data with log transformation was used to suitable for deep learning prediction with smaller magnitude variation as target data.

#### SRS imaging data

Data across all channels was normalized by RMS normalization, which was tested to have the best prediction performance compared with min-max normalization and simple linear unmixing. The across channel images for the same tissue sample are manually aligned to minimize location shifts.

#### Image registration

Consider the scale mismatch between MALDI and FTIR/SRS datasets, MALDI data was first upscaled to match the spatial scale of FTIR or SRS datasets. Then we followed the protocol of reported method [7], which performs rough alignment first with manually picked registration markers using affine transformation, and then conducts intensity-based refined registration. MALDI dataset was set as the standard and the same transformation matrix was applied across all channels for either FTIR data or SRS data. The registration result is provided in Supplementary Fig. 14.

## Data analysis

Both CCA analysis and regression analysis were performed using Python Scikit-learn. For CCA, the model was fitted to the entire dataset of matched pixels from IR (X) and MALDI (Y) measurements, with two components (n_components=2) without scaling (scale=False) to maintain the natural variance structure of the data. This generated two pairs of canonical variates: (U₁, V₁) and (U₂, V₂), representing the first and second canonical components for IR and MALDI data, respectively. For regression analysis, the transformation matrix was built on one paired whole-brain sagittal brain data for MALDI and FTIR.

### Modeling

#### Partial least square (PLS) regression

PLS was performed using Python Scikit-learn. 3D hyperspectral imaging data was reshaped into 2D with spectrum peaks as features. Each whole-brain sagittal section data contains around 8 million spectra. PLS model was trained on either 4 pairs of whole-brain sagittal section data reshaped in 2D format, where FTIR data consists of 207 spectrum features and MALDI ground truth data contains 100 targets. Hyperparameters were finetuned via cross-validation, with 10 number of components kept. One PLS model was trained to predict 100 m/z peaks in MALDI data.

### Deep learning

Our Hyper-pix2pix model used pix2pix framework with further modification for hyperspectral image translation. U-Net model is used as the image generator/translator. Its encoder progressively downsamples the input image using convolutional layers, with 4 sequential blocks and each block as: Conv2D (stride 2, kernel size 4, padding 1), GroupNorm2D, LeakyReLU. The first layer is a Conv2d layer to convert input SRS input channels to designed input channels for down-sampling operation. The decoder contains 4 upsampling blocks, with each block as: ConvTranspose2D (stride 2, kernel size 4, padding 1), GroupNorm2D, ReLU. A softplus operation is added in the last layer to ensure non-negative predictions, matching the ground truth. For discriminator, we adopted the PatchGAN discriminator with 3 blocks, with each block as: Conv2D, BatchNorm2D, LeakyReLU. Note that noise was not added in input data to ensure discriminate outputs from U-Net generator/translator.

We designed a loss function tailored for predictions across hyperspectral channels. The loss function contains three parts, including cGAN loss, L2 loss, and spectrum loss, which is spectrum angle mapping (SAM) loss to enforce similar spectrum profile between predictions and targets. The SAM was calculated by reshaping 3D image data into 2D. Although L1 loss was suggested by several papers in image translation to reduce blurry outputs, we found L2 loss works better in this task to translate SRS images into MALDI images. In addition, we tested multiple loss functions and combinations, including Charnonnier loss, correlation loss, etc. The current loss function provides the best prediction results. In the equations below, x represents SRS input data, y represents MALDI targets, G and D represents generator and discriminator, respectively.

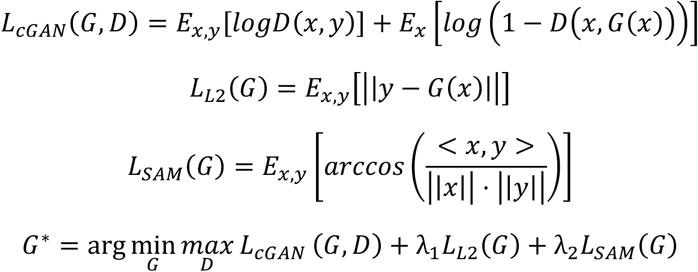

#### Ensemble learning

Considering the large size of SRS images for mouse-brain quarter coronal section (∼ 5000 x 3000) per image, the input datasets were divided into multi-scale patches, including 256 x 256, 512 x 512 and 1024 x 1024. To capture spatial features across different scales, one model was trained for each scale of image patches. The final outputs are the weighted sum of three models trained given input at different scales, with coefficient as 0.4, 0.4, 0.2 for models trained on 256, 512 and 1024 patches, respectively.

#### Model training

The designed deep learning models were trained on a single NVIDIA 4090 GPU. Data augmentation, including rotation, horizontal flip and vertical flip, was used to improve dataset size. Adam optimizer was used for both G and D with starting rate as 1e-4, and learning rate decay over epochs. D and G were alternatively trained. Batch size was selected as 64 for 256 x 256 inputs, 16 for 512 x 512 inputs, and 4 for 1024 x 1024 inputs. 200 epochs were trained for each model.

#### Evaluation

PSNR, SSIM and PCC are used as the three metrics for evaluation. For PLS prediction, MALDI data is kept in its original scale for both training and evaluation considering the high correlation between two datasets at the original scale. For deep learning, MALDI data is log transformed (np.log(1+x)) to reduce magnitude variation in targets in model training, which was found to have better performance than directly using the original scale. For evaluation, both the prediction and target data are mapped back to the original scale for comparative evaluation regrading PLS.

### Theory

For one pixel in IR imaging dataset its IR absorbance at a particular wavelength λ (*E*_n,λ_) can be explained by Beer-lambert law as shown in equation 6, where *c*_9_ represents concentration of a specific biological analyte, ℇ_9,8_ represents molar extinction coefficient of the this analyte at this particular wavelength, *l* is a constant representing the optical length at this pixel, and *n_c_* represents the total number of biological analyte categories (components). Accordingly, ***c_n_*** represents the concentration vector of all biological analytes and ℇ**_λ_** represents the corresponding molar extinction coefficient of each analyte at the given wavelength. This equation simply describes the pixel absorbance as a linear combination of all the biological analytes’ absorbance located at this pixel at a given wavelength. For the same pixel in MALDI imaging, the mass spectrum intensity at a particular m/z value (*M_n,m_*) can be described in equation 7, where *c*_j_′ represents a particular biological analyte and *r_j,m_* is a proportionality constant that describing the response of analyte *c_j_* at this m/z values. It includes factors such as the ionization efficiency and transmission efficiency that affect the signal. Note that due to the high discrimination power of mass spectrum, one m/z value generally corresponds to only one type of analyte.

When looking at the whole spectrum at a given pixel, the connection between two modalities becomes more clear. As shown in equation 8, The whole IR spectrum ***E_n_*** (a vector) can be expressed as the multiplication result between the concentration vector ***c_n_*** and the extinction coefficient matrix ℇ, times the constant l, where n_e_ represents the number of wavelengths. Accordingly, the whole mass spectrum can be expressed as the multiplication between mass response matrix **R_m_** and concentration vector ***c_n_***′, where n_m_ represents the number of masses in mass spectrum and ***n_c_***′ represents the number of biological analytes measured in mass-spec imaging.

When expanding to the whole hyperspectral imaging dataset, the connection between two modalities becomes clear. As shown in equation 10, the whole IR imaging data E is the matrix multiplication result between concentration matrix and extinction coefficient matrix ℇ constant l, where N represents the number of pixels in the imaging dataset. If let ***R_e_*** = *l*ℇ, then the final expression in equation 1 can be derived. Similarly, the whole MALDI imaging data can be expressed in equation 11.

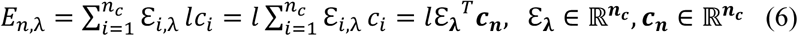

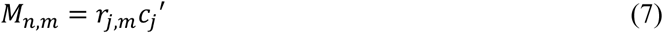

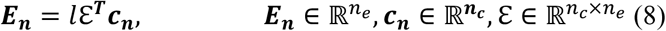

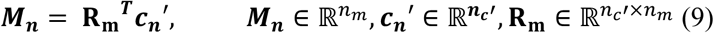

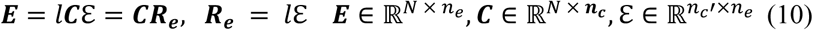

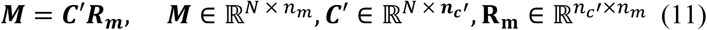

In equation (5), we can see that the unconnected parts between **E** and **M** are ***C_m_R_m_***_,***m***_ and ***C_e_R_e_***_,***e***_. If ***C_m_***, ***C_e_*** and **C_o_** are linearly dependent, it can be inferred that **M** can be fully expressed by **E**, when ignoring noises. An example is shown below (there are many cases for the linear dependencies and we show an ideal case here), where ***C_e_*** (e.g, proteins) is linearly combination of **C_o_** (amino acids and peptides), and Cm (e.g. choline) is also a linear combination of **C_o_** (fatty acids), then **M** can be fully expressed by **E** via linear operation. It can be easily inferred that if ***C_e_*** is linearly dependent by ***C_m_*, M** can be fully expressed by **E** via linear operation as well.

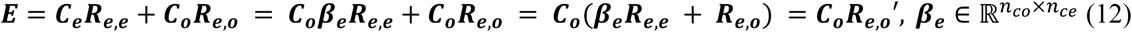

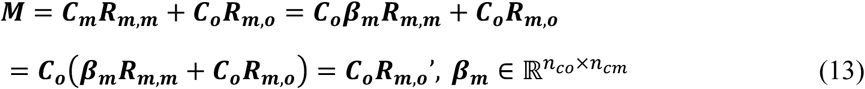

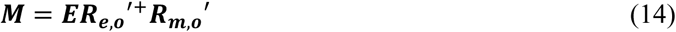

If the relation of ***C_m_***, ***C_e_*** and **C_o_** are not linearly dependent, M can still be expressed by E but through a more complex format. An example is show below, where ***f_e_*** is function describing the association between ***C_o_*** and ***C_e_***, and where ***f_m_*** is function describing the association between ***C_o_*** and ***C_m_***. If ***f_e_*** and ***f_m_*** are revertible, **M** can be expressed by **E** in equation 17. Although the specific format of are unknown, deep learning models can be used to approximate these functions.

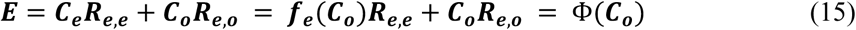

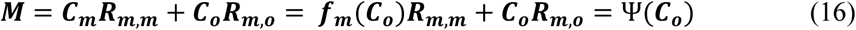

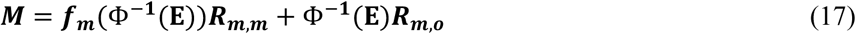

## Statistical analysis

For the CCA analysis, we analyzed matched IR and MALDI imaging datasets from mouse sagittal whole-brain tissue. Initial data dimensions were 3975 × 2220 × 207 for IR measurements and 3975 × 2220 × 100 for MALDI measurements. After remove background pixels, 5,083,282 valid tissue pixels were left for analysis. The final processed data matrices consisted of 5,083,282 matched spatial pixels with 207 IR spectral variables and 100 MALDI m/z peaks. CCA was applied with two components, yielding correlation coefficients of 0.914 and 0.742 for the first and second pairs of canonical components, respectively. Figure 2c-2d contains data points from 5,083,282 pixels. For the latent space visualization (Figure 2e), data was down-sampled to 100,000 points to ensure computational efficiency while maintaining representative distributions.

## Reference

1. Wei, L. et al. Super-multiplex vibrational imaging. Nature 544, 465–470 (2017).

2. Hu, F. et al. Supermultiplexed optical imaging and barcoding with engineered polyynes. Nat. Methods 15, 194–200 (2018).

3. Shi, L. et al. Highly-multiplexed volumetric mapping with Raman dye imaging and tissue clearing. Nat. Biotechnol. 40, 364–373 (2022).

4. Gerdes, M. J. et al. Highly multiplexed single-cell analysis of formalin-fixed, paraffin-embedded cancer tissue. Proc. Natl. Acad. Sci. U. S. A. 110, 11982–11987 (2013).

5. Lin, J.-R. et al. Highly multiplexed immunofluorescence imaging of human tissues and tumors using t-CyCIF and conventional optical microscopes. Elife 7, (2018).

6. Gut, G., Herrmann, M. D. & Pelkmans, L. Multiplexed protein maps link subcellular organization to cellular states. Science 361, eaar7042 (2018).

7. Chen, K. H., Boettiger, A. N., Moffitt, J. R., Wang, S. & Zhuang, X. RNA imaging. Spatially resolved, highly multiplexed RNA profiling in single cells. Science 348, aaa6090 (2015).

8. Goltsev, Y. et al. Deep profiling of mouse splenic architecture with CODEX multiplexed imaging. Cell 174, 968–981.e15 (2018).

9. Eng, C.-H. L. et al. Transcriptome-scale super-resolved imaging in tissues by RNA seqFISH. Nature 568, 235–239 (2019).

10. Jang, C., Chen, L. & Rabinowitz, J. D. Metabolomics and isotope tracing. Cell 173, 822–837 (2018).

11. Bhargava, R. Infrared spectroscopic imaging: the next generation. Appl. Spectrosc. 66, 1091–1120 (2012).

12. Cheng, J.-X. & Xie, X. S. Vibrational spectroscopic imaging of living systems: An emerging platform for biology and medicine. Science 350, aaa8870 (2015).

13. Oh, S. et al. Protein and lipid mass concentration measurement in tissues by stimulated Raman scattering microscopy. Proc. Natl. Acad. Sci. U. S. A. 119, e2117938119 (2022).

14. Ho, C.-S. et al. Rapid identification of pathogenic bacteria using Raman spectroscopy and deep learning. Nat. Commun. 10, 4927 (2019).

15. Hollon, T. C. et al. Near real-time intraoperative brain tumor diagnosis using stimulated Raman histology and deep neural networks. Nat. Med. 26, 52–58 (2020).

16. Liu, X. et al. Towards mapping mouse metabolic tissue atlas by mid-infrared imaging with heavy water labeling. Adv. Sci. (Weinh.) 9, e2105437 (2022).

17. Bhargava, R. Digital histopathology by infrared spectroscopic imaging. Annu. Rev. Anal. Chem. (Palo Alto Calif.) 16, 205–230 (2023).

18. Liu, X., Shi, L., Zhao, Z., Shu, J. & Min, W. VIBRANT: spectral profiling for single-cell drug responses. Nat. Methods 21, 501–511 (2024).

19. Zhang, W. et al. Multi-molecular hyperspectral PRM-SRS microscopy. Nat. Commun. 15, 1599 (2024).

20. Liu, Z. et al. Virtual formalin-fixed and paraffin-embedded staining of fresh brain tissue via stimulated Raman CycleGAN model. Sci. Adv. 10, eadn3426 (2024).

21. Kondepudi, A. et al. Foundation models for fast, label-free detection of glioma infiltration. Nature 637, 439–445 (2025).

22. Buchberger, A. R., DeLaney, K., Johnson, J. & Li, L. Mass spectrometry imaging: A review of emerging advancements and future insights. Anal. Chem. 90, 240–265 (2018).

23. Rappez, L. et al. SpaceM reveals metabolic states of single cells. Nat. Methods 18, 799–805 (2021).

24. Norris, J. L. & Caprioli, R. M. Analysis of tissue specimens by matrix-assisted laser desorption/ionization imaging mass spectrometry in biological and clinical research. Chem. Rev. 113, 2309–2342 (2013).

25. Van de Plas, R., Yang, J., Spraggins, J. & Caprioli, R. M. Image fusion of mass spectrometry and microscopy: a multimodality paradigm for molecular tissue mapping. Nat. Methods 12, 366–372 (2015).

26. Hu, F., Shi, L. & Min, W. Biological imaging of chemical bonds by stimulated Raman scattering microscopy. Nat. Methods 16, 830–842 (2019).

27. Stimulated Raman Scattering Microscopy: Techniques and Applications. (Elsevier - Health Sciences Division, Philadelphia, PA, 2021).

28. Min, W., Cheng, J. X. & Ozeki, Y. Theory, innovations and applications of stimulated Raman scattering microscopy. Nat. Photonics in press (2025).

29. Baker, M. J. et al. Clinical applications of infrared and Raman spectroscopy: state of play and future challenges. Analyst 143, 1934–1934 (2018).

30. Felten, J. et al. Vibrational spectroscopic image analysis of biological material using multivariate curve resolution-alternating least squares (MCR-ALS). Nat. Protoc. 10, 217–240 (2015).

31. Palmer, A. et al. FDR-controlled metabolite annotation for high-resolution imaging mass spectrometry. Nat. Methods 14, 57–60 (2017).

32. Wadie, B. et al. METASPACE-ML: Context-specific metabolite annotation for imaging mass spectrometry using machine learning. Nat. Commun. 15, 9110 (2024).

33. Clarke, H. A. et al. Spatial mapping of the brain metabolome lipidome and glycome. Nat. Commun. 16, 4373 (2025).

34. Hardoon, D. R., Szedmak, S. & Shawe-Taylor, J. Canonical correlation analysis: an overview with application to learning methods. Neural Comput. 16, 2639–2664 (2004).

35. Geladi, P. & Kowalski, B. R. Partial least-squares regression: a tutorial. Anal. Chim. Acta 185, 1–17 (1986).

36. Wang, Z., Bovik, A. C., Sheikh, H. R. & Simoncelli, E. P. Image quality assessment: from error visibility to structural similarity. IEEE Trans. Image Process. 13, 600–612 (2004).

37. Zhang, D. et al. Inferring super-resolution tissue architecture by integrating spatial transcriptomics with histology. Nat. Biotechnol. 42, 1372–1377 (2024).

38. Jia, Y., Liu, J., Chen, L., Zhao, T. & Wang, Y. THItoGene: a deep learning method for predicting spatial transcriptomics from histological images. Brief. Bioinform. 25, (2023).

39. Schmauch, B. et al. A deep learning model to predict RNA-Seq expression of tumours from whole slide images. Nat. Commun. 11, 3877 (2020).

40. Min, W. & Gao, X. Absolute signal of stimulated Raman scattering microscopy: A quantum electrodynamics treatment. Sci. Adv. 10, eadm8424 (2024).

41. Isola, P., Zhu, J.-Y., Zhou, T. & Efros, A. A. Image-to-image translation with conditional adversarial networks. in 2017 IEEE Conference on Computer Vision and Pattern Recognition (CVPR) 5967–5976 (IEEE, 2017).

42. Ronneberger, O., Fischer, P. & Brox, T. U-Net: Convolutional Networks for Biomedical Image Segmentation. arXiv [cs.CV] (2015).

43. Manifold, B., Men, S., Hu, R. & Fu, D. A versatile deep learning architecture for classification and label-free prediction of hyperspectral images. *Nat*. Mach. Intell. 3, 306–315 (2021).

44. Ben-Israel, A. & Greville, T. N. E. Generalized Inverses: Theory and Applications. (Springer, New York, NY, 2003).

45. Levine, J. H. et al. Data-driven phenotypic dissection of AML reveals progenitor-like cells that correlate with prognosis. Cell 162, 184–197 (2015).

46. Rafelski, S. M. & Theriot, J. A. Establishing a conceptual framework for holistic cell states and state transitions. Cell 187, 2633–2651 (2024).

47. Rood, J. E., Hupalowska, A. & Regev, A. Toward a foundation model of causal cell and tissue biology with a Perturbation Cell and Tissue Atlas. Cell 187, 4520–4545 (2024).

48. Jumper, J. et al. Highly accurate protein structure prediction with AlphaFold. Nature 596, 583–589 (2021).

49. Abramson, J. et al. Accurate structure prediction of biomolecular interactions with AlphaFold 3. Nature 630, 493–500 (2024).

50. Saar, B. G. et al. Video-rate molecular imaging in vivo with stimulated Raman scattering. Science 330, 1368–1370 (2010).

51. Ozeki, Y. et al. High-speed molecular spectral imaging of tissue with stimulated Raman scattering. Nat. Photonics 6, 845–851 (2012).

52. Lin, H. et al. Microsecond fingerprint stimulated Raman spectroscopic imaging by ultrafast tuning and spatial-spectral learning. Nat. Commun. 12, 3052 (2021).

53. Lin, H. et al. Label-free nanoscopy of cell metabolism by ultrasensitive reweighted visible stimulated Raman scattering. Nat. Methods 1–11 (2025).

54. Bassan, P., Weida, M. J., Rowlette, J. & Gardner, P. Large scale infrared imaging of tissue micro arrays (TMAs) using a tunable Quantum Cascade Laser (QCL) based microscope. Analyst 139, 3856–3859 (2014).

55. Yeh, K., Kenkel, S., Liu, J.-N. & Bhargava, R. Fast infrared chemical imaging with a quantum cascade laser. Anal. Chem. 87, 485–493 (2015).

56. Bird, B. & Baker, M. J. Quantum cascade lasers in biomedical infrared imaging. Trends Biotechnol. 33, 557–558 (2015).

57. Orringer, D. A. et al. Rapid intraoperative histology of unprocessed surgical specimens via fibre-laser-based stimulated Raman scattering microscopy. *Nat*. Biomed. Eng. 1, 0027 (2017).

58. Smolina, M. & Goormaghtigh, E. Gene expression data and FTIR spectra provide a similar phenotypic description of breast cancer cell lines in 2D and 3D cultures. Analyst 143, 2520–2530 (2018).

59. Kobayashi-Kirschvink, K. J. et al. Prediction of single-cell RNA expression profiles in live cells by Raman microscopy with Raman2RNA. Nat. Biotechnol. 42, 1726–1734 (2024).

## Methods Reference

1. Veerasammy, K. et al. Sample preparation for metabolic profiling using MALDI mass spectrometry imaging. J. Vis. Exp. (2020) doi:10.3791/62008.

2. Ding, J. et al. A metabolome atlas of the aging mouse brain. Nat. Commun. 12, 6021 (2021).

3. Wang, J. et al. MALDI-TOF MS imaging of metabolites with a N-(1-naphthyl) ethylenediamine dihydrochloride matrix and its application to colorectal cancer liver metastasis. Anal. Chem. 87, 422–430 (2015).

4. Khamidova, N. et al. DBDA matrix increases ion abundance of fatty acids and sulfatides in MALDI-TOF and mass spectrometry imaging studies. J. Am. Soc. Mass Spectrom. 34, 1593–1597 (2023).

5. Liu, Y. et al. Integrated mass spectrometry imaging reveals spatial-metabolic alteration in diabetic cardiomyopathy and the intervention effects of ferulic acid. J. Pharm. Anal. 13, 1496–1509 (2023).

6. Behdenna, A. et al. pyComBat, a Python tool for batch effects correction in high-throughput molecular data using empirical Bayes methods. BMC Bioinformatics 24, 459 (2023)

7. Neumann, E. K. et al. Multimodal chemical analysis of the brain by high mass resolution mass spectrometry and infrared spectroscopic imaging. Anal. Chem. 90, 11572–11580 (2018).

8. Smith, K. W., Fecke, A., Kasarla, S. S. & Phapale, P. Large-scale metabolite imaging gallery of mouse organ tissues to study spatial metabolism. J. Proteome Res. 24, 3105–3116 (2025).

9. Zhang, H. et al. TEMI: tissue-expansion mass-spectrometry imaging. Nat. Methods 22, 1051–1058 (2025).

